# The role of vertical and horizontal transmission in the assembly of seed fungal endophyte communities in wheat and wheat wild relatives

**DOI:** 10.1101/2022.11.21.517337

**Authors:** Or Sharon, Xiang Sun, Smadar Ezrati, Naomi Kagan-Trushina, Amir Sharon

## Abstract

Plants acquire fungal endophytes either from the environment or from their progenitors. These transmission modes are central in shaping the community as they affect species composition and balance. We studied fungal endophyte communities (FEC) and their seed-to-seed transmission in three Triticeae plant species: bread wheat (Triticum aestivum), wild emmer wheat (Triticum turgidum dicoccoides) and wild barley (Hordeum spontaneum). The FECs in the three plant species contained similar fungal taxa, however they were overall different. The most prevalent class of fungi was Dothideomycetes, which was dominated by the taxon Alternaria infectoria. In field collected plants, the number of taxa in the seeds was less than half the number in stems, with close to 90% of the taxa found in seeds also found in stems. Growing the same plant species in a controlled environment infection greatly affected their FEC composition; the FECs in the stems and seeds of these plants were richer and more diverse than in the original seeds, they were not dominated by a single taxon, and FECs in the new seeds had a similar richness and diversity to the stem FECs, with only 40% overlap. The controlled environment experiment confirmed vertical transmission of certain species, but also showed that external infection of the seeds is the main source for specific taxa, including A. infectoria. Collectively, our results show that many taxa can reach the seeds through the internal pathway, albeit in different abundance, and both internal and external sources significantly affect the composition of seed FECs.

## Introduction

All plant species contain fungal endophyte communities (FECs), which are part of their microbiome (Chen et al. 2021). These FECs are dynamic, they vary in composition between species and tissues, and are affected by plant developmental stage and environmental conditions. The number of endophytic fungal species within a given plant varies considerably, with recent studies indicating that hundreds of different amplicon sequence variants (ASVs) can be detected in some plant species (Gdanetz et al. 2021; de Souza et al. 2016; Fonseca-García et al. 2016). Certain fungal endophytes have well-established beneficial effects on their hosts, such as protection from pathogens, improved performance under abiotic stress, and growth promotion (Llorens et al. 2019; Bouzouina et al. 2021; Bastias et al. 2017; Chitnis et al. 2020; Chen et al. 2021). However, the role of the bulk of the species and their relationships with the host plants remain unclear.

Plants can acquire fungal endophytes in two ways: internally, from the maternal plant to the offspring through the seed (known as vertical transmission), and externally, from the soil, water, or air (known as horizontal transmission). Vertical transmission has generally been considered rare and restricted to specific fungal taxa, such as certain Epichloë species (Liu et al. 2017), but recent studies indicates that it is likely more widespread and includes many more species than previously assumed (Hodgson et al. 2014; Abdelfattah et al. 2021; Özkurt et al. 2020; Fort et al. 2021).

Endophytes can reach the new seeds internally (vertical transmission), by growing in the xylem of the progenitor plant, or externally (horizontal transmission), by growing through the pollen of the progenitor plant or through external contamination of the seed (Barret et al. 2015). Vertically transmitted (internal pathway) endophytes can reach the embryo (Liu et al. 2017), while the external pathways are thought to lead to infection of the endosperm or the seed coat (Barret et al. 2016). Seeds will thus contain two distinct fungal populations: the first represents the true seed-borne endophytes, and the second represents other seed-residing species (internal seed coat and endosperm) that originate either from the internal or external pathway (through the pollen or direct seed infection). As it is difficult to distinguish between the two populations, most studies refer to them collectively as a single seed FEC (Özkurt et al. 2020; Morales Moreira et al. 2021; Simonin et al. 2022; Rochefort et al. 2021). Determining the mode of transmission can also be technically challenging due to a fungal species overlap between seed FECs and soil or air FECs (Shade et al. 2017). Whether acquired internally or externally, seed-borne endophytes are the most immediate source of inoculum in seedlings and can thus significantly affect both plant development and the FEC structure within the plant (Nelson 2018). The vertically-transmitted taxa represent a stable portion of the community, and it is logical to assume that they are more important for plant performance than occasional taxa acquired from the environment (Christian et al. 2017). Hence, discriminating between the vertically- and horizontally transmitted taxa can help further understand the mechanisms that shape the community and aid identification of beneficial taxa.

Studies on endophytes in wheat and related species have provided a wealth of information on the richness and composition of FECs in below- and above-ground plant parts (Kinnunen-Grubb et al. 2020; Chen et al. 2019; Hassani et al. 2020; Rojas et al. 2020; Latz et al. 2021; Sun et al. 2020a, 2020b). These studies show that various factors such as plant genotype, domestication, and environmental influences affect the composition of the community to some degree. Collectively, these studies show that the bulk of the population is highly sporadic. However, there is also a set of core taxa found in all these studies, including genera of filamentous ascomycetes, e.g., *Alternaria* and *Cladosporium*, and basidiomycete yeasts, e.g., *Vishniacozyma* and *Sporobolomyces*. Studies on the origin of the fungi and interrelationships between the populations within the plant that focus on seed-to-seed transmission are scarce and more research is necessary to clarify the true nature of seeds fungal endophytes.

Here, we examined seed endophytes populations in three Triticeae species: bread wheat (*Triticum aestivum* L; *Ta*), wild emmer wheat (*Triticum turgidum dicoccoides*; *Td*) and wild barley (*Hordeum spontaneum*; *Hs*). We show that while the seed FECs in the three plant species share most of their taxa and are dominated by *Alternaria infectoria*, the FECs in the wild plants are richer and more diverse than the bread wheat FECs. We also showed that while the most dominant taxa can reach the seeds internally, external seed infection is the primary source of certain taxa, including *A. infectoria*. Finally, the composition of seed and stem FECs in greenhouse-grown plants differs dramatically to that of field-collected plants, suggesting that environmental conditions play central roles both in shaping the community, and in facilitating the exchange of endophytes between stems to seeds.

## Results

### Structure of the seed FECs in the three plant species

Seeds of *Ta, Td*, and *Hs* were collected from mature plants growing in agricultural fields (*Ta*) and natural habitats (*Td, Hs*) (Supplementary Table S1). To characterize the seed FECs, we extracted DNA and performed ITS amplicon sequencing of 420 seed samples. After filtering and quality control, we obtained a final dataset that included 62 taxa in 382 samples (Supplementary Table S2), with a total of 6,548,757 reads (Supplementary Table S3).

Alpha diversity observed taxa and Shannon diversity index showed that FECs from *Hs* were the most diverse, with the FECs in *Hs* and *Td* exhibiting higher diversity than the *Ta* FEC (Fig. 1A, Supplementary Table S4). Analysis of beta diversity showed significant differences between seed FECs of all three plant species. Principal coordinates analysis (PCoA) of abundance using Bray–Curits dissimilarities revealed an overlap between samples from different plant species (Fig. 1B). However, the communities were statistically different according to PERMANOVA analysis (F = 18.93, R^2^ = 0.09, p < 0.001), and pairwise adonis results were significant (*P* = 0.001) for all plant species combination (Supplementary Table S5). These results are in agreement with a previous study showing that diversity of stem FECs from wild plant species was higher than the diversity of stem FECs from bread wheat (Sun et al. 2020b). The entire seed FEC contained 62 taxa, 44 (71%) of them in the *Ascomycota* and the rest in the *Basidiomycota*. Among the 62 taxa, 39 were common to all three plant species, nine taxa were species-specific, and the rest were common to two of the three plant species (Fig. 1C). About one third of the taxa were yeast or yeast-like, with most of them in the phylum *Basidiomycota*, meaning that most of the basidiomycetes taxa were yeasts. *Dothideomycetes* was the most dominant class in the entire community (50% of the taxa), followed by *Tremellomycetes* (basidiomycete yeasts, 19%). Despite some differences in relative abundance between the plant species, the top 20 most abundant and prevalent taxa were found in all plant hosts. Taxa belonging to the family *Pleosporaceae* were the most abundant in all plant species and were dominated by *Alternaria infectoria*, which was found in close to 100% of the samples and had a relative abundance of 0.68 in the entire FEC (Supplementary Table 6). The next most abundant taxa were *Alternaria alternata, Cladosporium cladosporioides* and *Mycosphaerella tassiana*, with some variation in abundance between plant species. For example, *Chalastospora gossypii* was the third most abundant fungal taxon in *Td*, while in *Ta, Stemphylium ASV6* was the second most abundant taxon after *A. infectoria*, and *C. cladosporioides* was only eighth. It can be concluded that the seed FECs in the different plant species contain a similar set of taxa that is dominated by *A. infectoria*. However, differences do exist in the structure of the communities, which can be attributed to the plant species.

**Fig 1.**
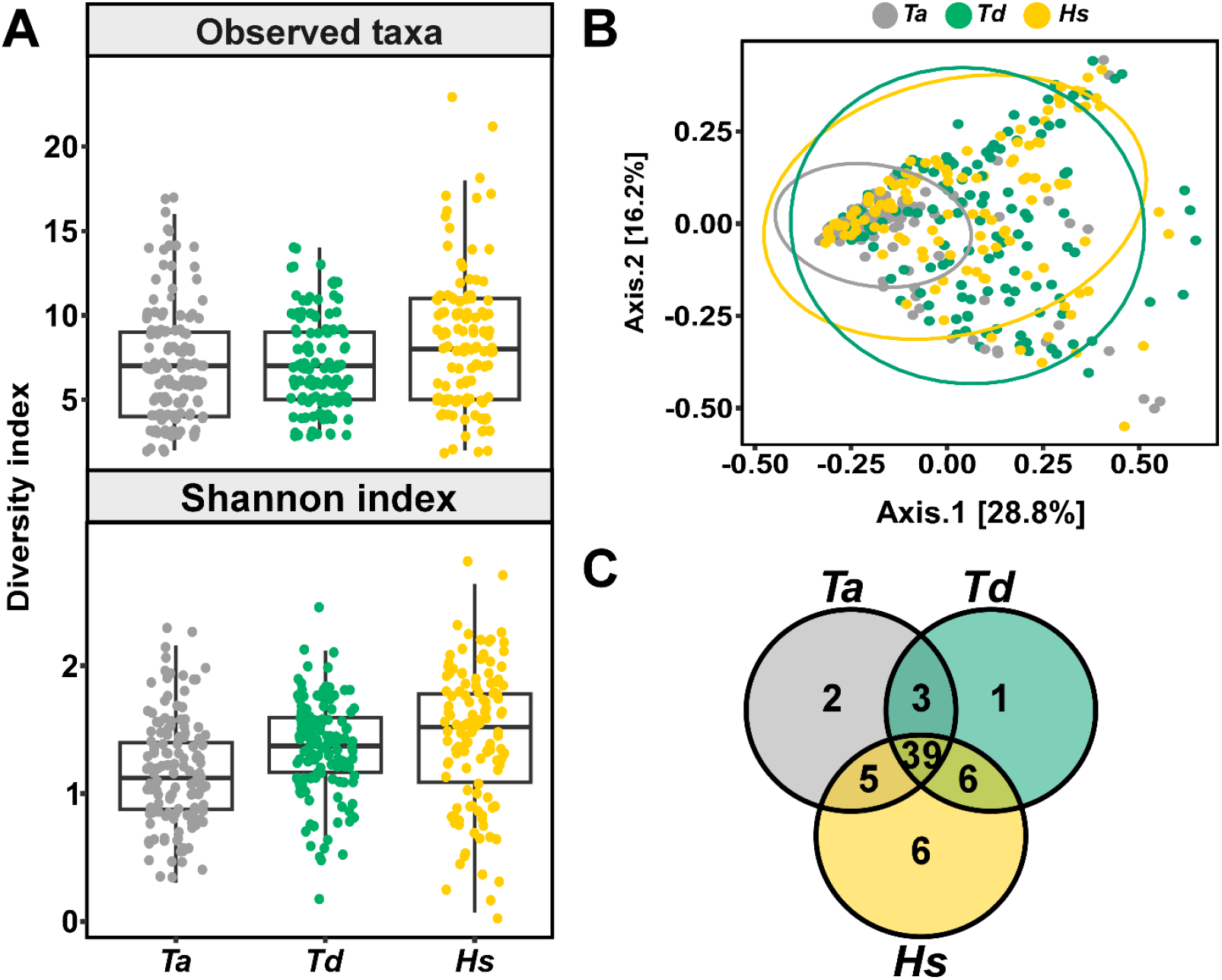
Composition of FECs in field-collected seeds. Seeds of bread wheat (*Ta*), wild wheat (*Td*), and wild barley (*Hs*) were sampled from cultivated seeds (*Ta*) and natural habitats (*Td* and *Hs*). DNA was extracted from 160 samples of each plant species and ITS amplicons were produced using the ITS1F-ITS2 fungal universal primers. (A) Alpha diversity parameters of the FECs in each plant species. Top – observed, bottom - Shannon index. Each dot in the boxplot represents the diversity in one sample. The communities were found to be statistically different (*P* < 0.001) according to Kruskal–Wallis test followed by pairwise Wilcox test, see supplementary table S4 for details. (B) Principal coordinate analysis (PCoA) of the seeds FECs based on Bray-Curtis dissimilarity matrix using Hellinger transformed abundance data. The three FECs were found to be statistically different according to PERMANOVA analysis. (C) Venn diagram of the fungal taxa in seeds of the different plant species. Color code in all panels: *Ta* – gray, *Td* - green, *Hs* - yellow.

### Stems FECs are more diverse than seed FECs

To determine the association between the seed and stem FECs, we analyzed ITS amplicon sequencing from the DNA of previously collected stems from the three plant species (Sun et al. 2020a, 2020b) together with amplicons from seeds. Following filtering and quality control, the entire dataset contained 735 samples (382 seeds and 353 stems), which produced 9,601,707 reads and 250 taxa (Supplementary Table S2 and S3). The average read number in stem samples was 8,648, about half the number of reads (17,143) in the seed samples, but the number of taxa in the stems was on average 2.6 times that of seeds, in all plant hosts and in each of the plant species. Likewise, for all plant species the alpha diversity of the stem FECs was significantly higher than in seeds (Fig. 2A); on average, the FECs from stems were twice more diverse than in seeds, and in *Td* stems the Shannon index was more than 3 times higher than in seeds (Supplementary Table S8). The higher diversity of stem FECs was reflected by the different composition of the seed and stem FECs as measured by PCoA, and PERMANOVA analysis of Bray–Curits dissimilarities (F = 187.273, R^2^ = 0.19, *P* < 0.001 from the entire community) (Fig. 2B, Supplementary Table S9). Among the three plant species, the FECs from *Ta* seeds and stems formed the two most distinct clusters (F = 100.98, R^2^ = 0.27, *P* < 0.001). The dominant taxon in *Ta* stems was *Candida sake*, whereas the seeds of *Ta* were dominated by *A. infectoria*. By contrast, in *Td* and *Hs, A. infectoria* dominated the FECs in both the seeds and stems. This difference accounts at least in part for the greater separation between the stems and seeds FECs in *Ta* compared with the two other plant species.

**Fig 2.**
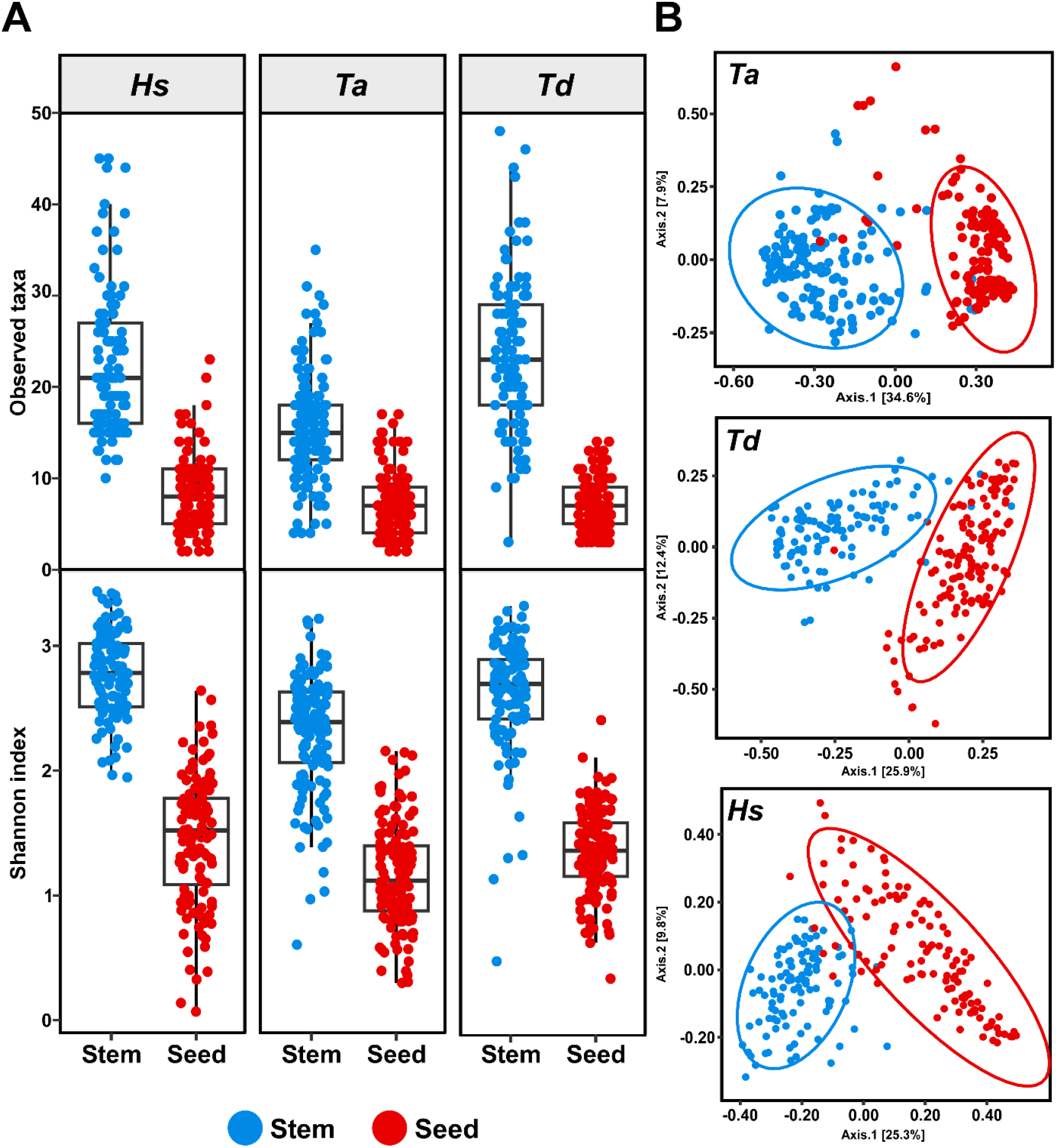
Comparative analysis of FECs in field-collected stems and seeds. (A) Alpha diversity: observed taxa (top) and Shannon diversity index (bottom). Each dot in the boxplot represents the diversity in one stem (blue) or seed (red) sample. Stems and seeds were statistically different (*P* < 0.001) according to Wilcox test (see supplementary table S8 for statistical details). (B) PCoA based on Bray-Curtis dissimilarity matrix. The stem and seed FECS are statistically different (*P* < 0.001) according to PERMANOVA using adonis analysis.

### Correlation between taxa in seed and stem FECs

The seeds of all three plant species had a lower number of taxa compared with stems (Fig. 3A). Importantly, apart from seven low abundance taxa, all the other 55 seed taxa were also found in stems, suggesting the stems as a main source of seed endophytes. We also noted a significant shift in the composition of the seed and stem FECs; *Dothideomycetes* was the most abundant class in the seed endophyte community (0.97) and accounted for 50% of the taxa present in the seed FECs (Fig. 3B). *Dothideomycetes* species were also the most prevalent class in stem FECs, but they were less dominant than in seeds, with 34% of the taxa and 0.43 abundance. Another significant difference was the lower abundance of yeast species in the seeds. Overall, the relative abundance of *Tremellomycetes* and *Saccharomycetes* yeasts was 0.2 and 0.14 in stems, but considerably lower in seeds at 0.007 and 0.002 respectively (Fig. 4). The most striking example of this is *C. sake*, the most prevalent yeast species in stems with a relative mean abundance of 0.28 and 86% incidence which dropped to only 0.004 abundance and less than 10% incidence in seeds (Supplementary Table S6 and S7). Contrary to yeasts, many of the filamentous taxa found in seeds were enriched compared to stems. For example, the relative abundance of *A. infectoria* increased from 0.3 in stems to 0.68 in seeds, and other *Alternaria* species, primarily *Alternaria alternata* and *Alternaria metachromatica*, exhibited a similar trend. A small number of filamentous taxa, in particular *Cladosporium* species, showed an opposite trend, possibly reflecting the slow growth rate of these species (Bullerman 2003).

**Fig 3.**
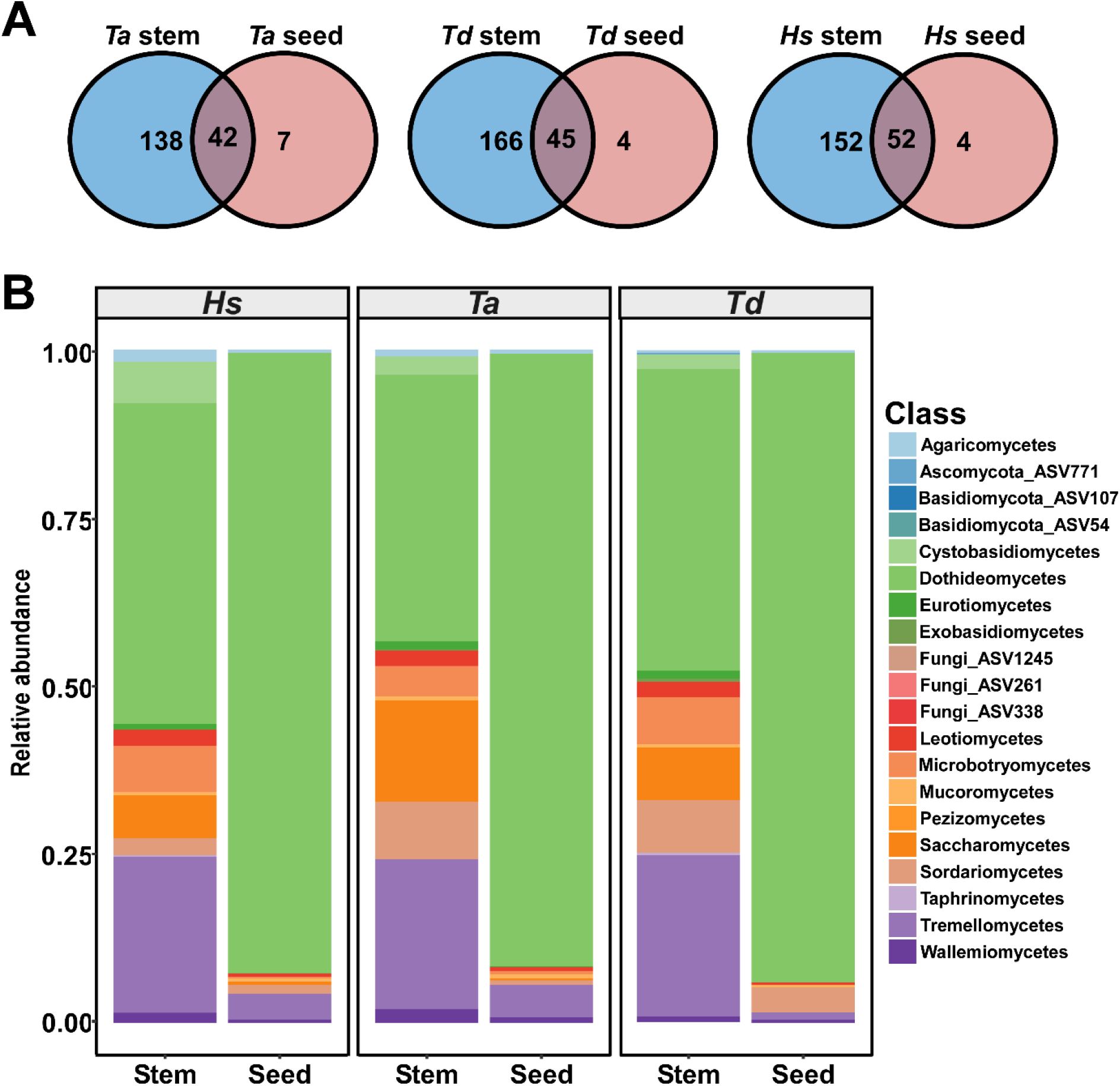
Distribution of taxa across plant species and organs in field-collected plants. (A) Venn diagram illustrating common and unique fungal taxa among stems and seeds in each bread wheat (*Ta*), wild wheat (*Td*), and wild barley (*Hs*). (B) Class-level relative abundance of fungal taxa in each of the six FECs.

**Fig 4.**
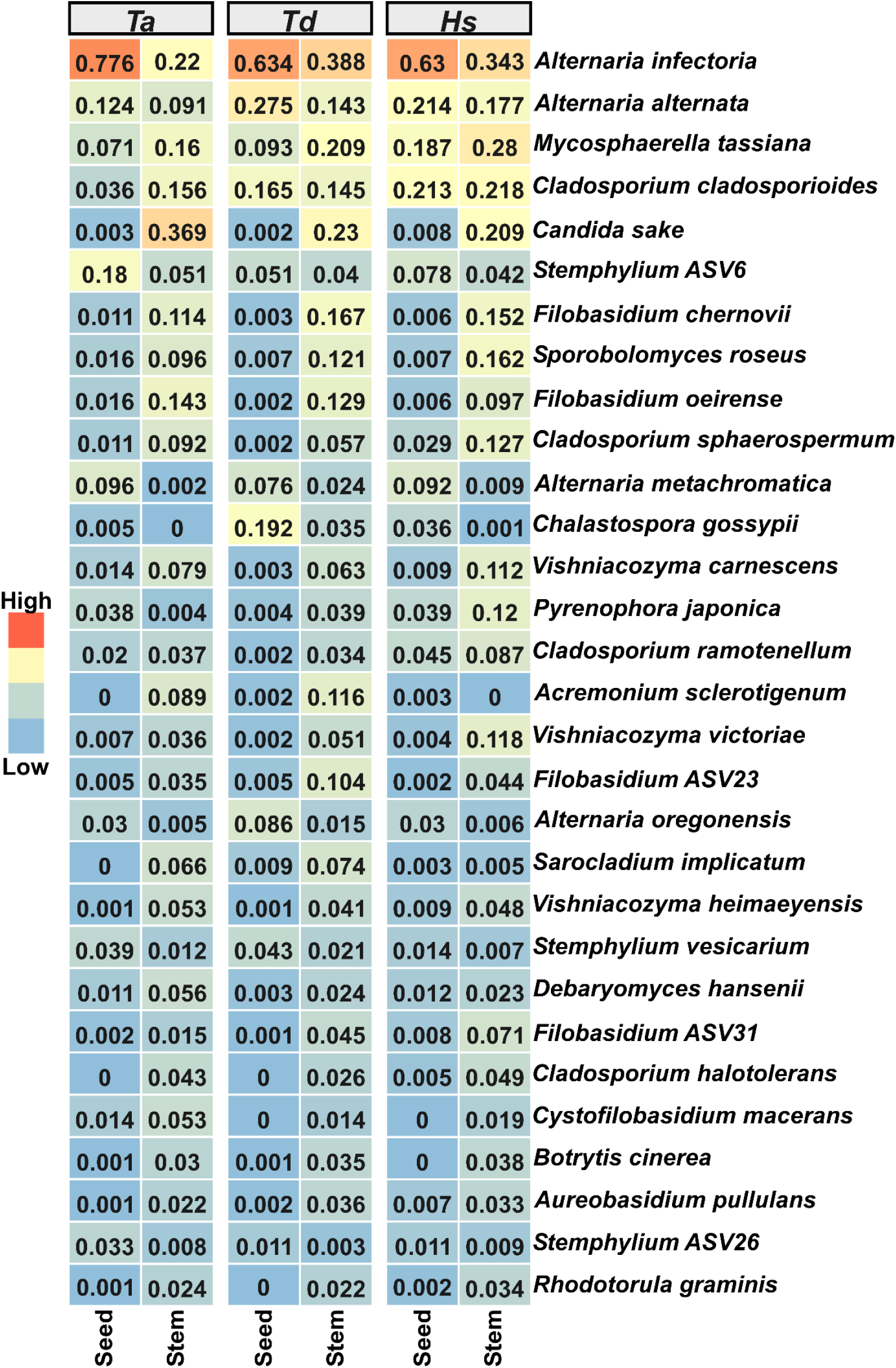
Relative abundance of the top 30 most abundant stem and seed taxa in field-collected samples. Values for each taxon represent the mean relative abundance.

### Seed-to-seed transmission of FECs under greenhouse conditions

The increased abundance of certain stem endophytes in seeds and the high overlap of the seed FEC with a portion of the stem community indicates that a significant part of the seed community originates in the stems. However, seeds of field-grown plants can also acquire endophytes from the environment (Barret et al. 2015) and therefore this source needs to be eliminated to determine the route and transmission mode of specific taxa. To address these issues and assess the significance of systemically transmitted endophytes, we produced plants in a greenhouse under controlled conditions from seeds that were collected in the wild (*Td* and *Hs*) or cultivated fields (*Ta*). To minimize external infection, we used autoclaved soil, and covered the spikes before anthesis with paper bags. We extracted DNA from soil samples, original seeds, stems that were at a similar developmental stage to stems in the field, and from new dry seeds, and performed ITS amplicon sequencing (Supplementary Table S10). Following QC and removal of samples with less than 1,000 reads, we obtained 27 sets of plant samples and 15 soil samples, with a total number of 1,772,592 reads yielding 365 taxa (Supplementary Table S11 and S12).

Strikingly, the FECs from stems and new seeds were greatly enriched compared with the original seeds; only 35 out of the 365 taxa were found in the original seeds, while 178 taxa were found in the stems and 202 in the new seeds. Out of the 35 seed taxa, 21 were also found in stems and 21 in the new seeds (Fig. 5A). The soil contained 83 taxa, 27 of them found in the new plants and the rest unique to the soil community. In accordance with the higher number of taxa, the FECs in the stems and seeds of greenhouse-grown plants were also more diverse than in the original seeds (Fig. 5B, Supplementary Table S13), and analysis of beta diversity showed clear separation between the different FECs (Fig. 5C). PREMANOVA (F= 10.77, R^2^= 0.248, p < 0.001) and pairwise adonis analyses showed statistical differences between all four FECs, as well as between the FECs in different plant species, with the exception of *Ta* and *Td* (Supplementary Table S14).

**Fig 5.**
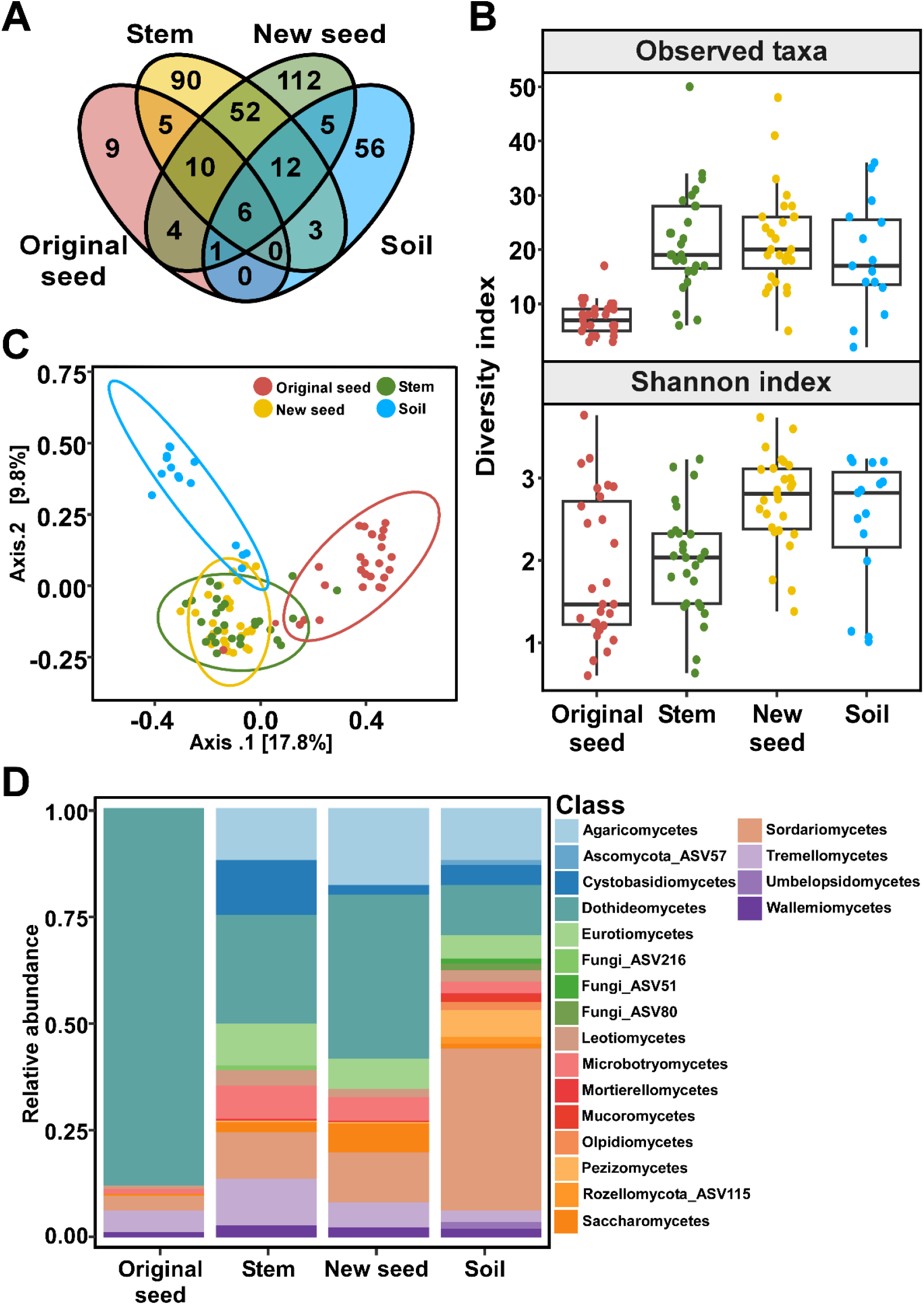
Composition and diversity of FECs in soil, original seeds, stem, and new seeds of greenhouse-grown plants. (A) Venn diagram of taxa in the original seeds, stems, new seeds, and soil. (B) Alpha diversity within sample types. Each dot in the boxplot represents the diversity in one sample. Original seeds differed from the rest of the samples. Significant differences were determined using Kruskal–Wallis test followed by pairwise Wilcox test (*P* < 0.001) (see supplementary table S13 for details). (C) PCoA of the FECs in soil, original seeds, stems, and new seeds based on Bray–Curtis dissimilarity. All FECs were statistically different from one another (*P* < 0.001) according to PERMANOVA using adonis analysis. (D) Relative abundance of the top 20 fungal classes across sample types.

### Changes in composition of FECs in greenhouse-grown plants compared with field-collected plants

Seeds and stems from the greenhouse-grown plants contained a similar number of taxa, but only 40% (80 taxa) of the seed taxa were also found in stems (Fig 5A). This contrasts with the situation in field-collected plants, in which the number of taxa in seeds was much smaller (25%) than in stems and 90% of the seed taxa were also found in the stems (Fig. 3A). Additionally, the composition of FECs in the greenhouse plants differed to that in field plants: the communities were more diverse, and there was no single highly dominant taxon. The highest number of taxa in the field-collected plants were in the class *Dothideomycetes* (64%, 0.96 relative abundance in the original seeds), whereas in the greenhouse plants (seeds and stems combined) the highest number of taxa belonged to the class *Agaricomycetes*. Surprisingly, however, we did not detect any *Agaricomycetes* taxa in the original seeds (Supplementary Table S15). Despite the significant reduction in the number of taxa, *Dothideomycetes* remained the most abundant taxa also in the greenhouse-grown plants, with 0.32 and 0.55 relative abundance in stems and seeds, respectively (Fig 5D). At the species level, the most significant difference between greenhouse- and field-grown plants was reduced abundance of *A. infectoria* (from 0.56 relative abundance in the original seeds to 0.05 and 0.13 in stems and seeds of greenhouse plants) and increased abundance of *Cladosporium* sp. (Supplementary Table S15 and S16). Hence, the main source of *A. infectoria*, and probably of other *Alternaria* species, in seeds seems to be external infection rather than vertical transmission. In other taxa, such as *Cladosporium* sp., the amounts in stems and seeds were similar in greenhouse- and field-grown plants, suggesting that the internal pathways are the main source of these species in the seeds. The most abundant *Cladosporium* species were *C. sphaerospermum* and *C. halotolerans*, with a combined relative abundance of 0.56 in the new seeds. Both were present in the autoclaved soil, which might have contributed to the increased abundance of these taxa in the greenhouse plants. *C. sake*, the most abundant yeast taxon in stems of field-collected plants, was not detected in the greenhouse plants. *F. chernovii* and *Vishniacozyma carnescens*, the two other most abundant yeast taxa in field-collected plants, were found in both stems and seeds of greenhouse plants, albeit at a relatively low abundance, suggesting that these species can be vertically transmitted.

The low overlap between the taxa found in the soil and original seeds and the stems and seeds of the greenhouse plants (only 16 shared taxa) is puzzling, since under the experimental conditions the soil and the original seeds are the only possible source of fungi. One explanation is that the lack of dominant taxa in the stems and new seeds of greenhouse-grown plants allows exposure of low abundant taxa that are otherwise not detected due to limited sequencing depth. In the original seeds, the top 10 most abundant taxa accounted for 83% of the total abundance and *A. infectoria* alone was responsible for 30% of that abundance, whereas in stems and seeds of greenhouse-grown plants the total abundance of the top 10 taxa was less than 40% (Fig 6). Moreover, there was no single dominant taxon in the greenhouse plants and the most abundant taxon in both stems (*Cystobasidium lysinophilum*) and new seeds (*C. sphaerospermum*) accounted for less than 10% of the total abundance. Of the 254 taxa unique to stems and new seeds, 86 were also found in samples collected from the field, suggesting that they were present in the original seeds but below detection threshold.

**Fig 6.**
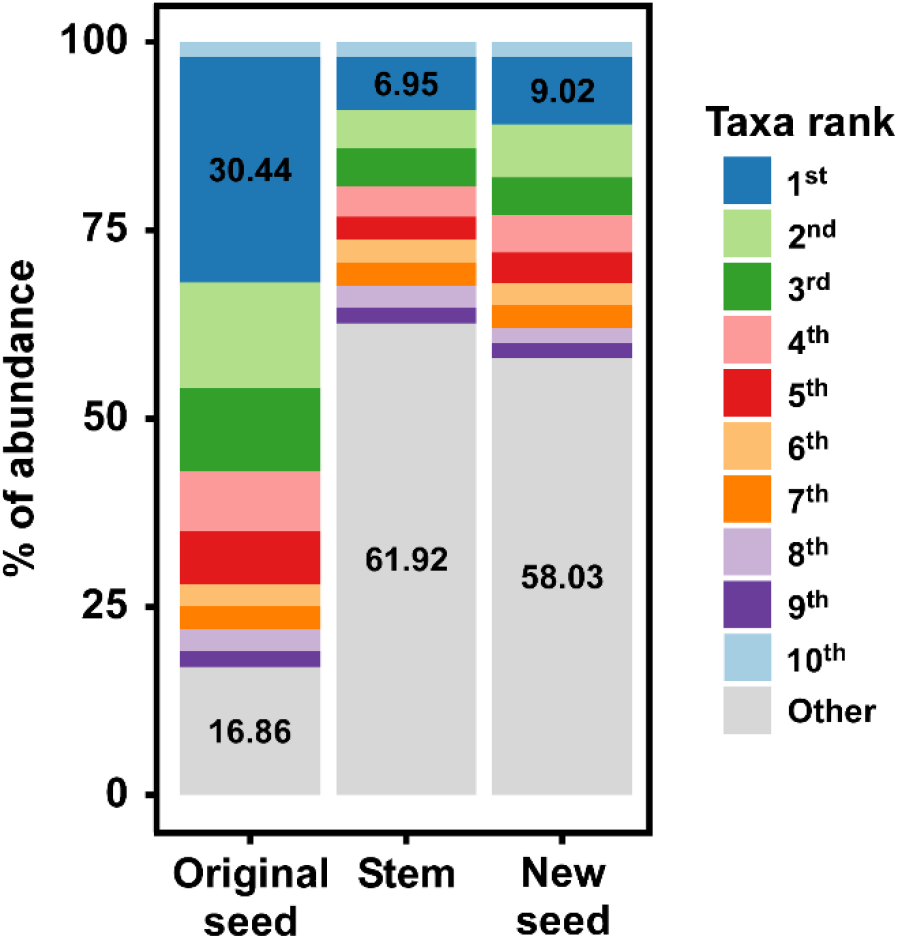
Abundance distribution of the top 10 most abundant taxa in original seeds, greenhouse-grown stems, and new seeds. Values were calculated by dividing the Hillinger transformed average abundance of each taxon by the sum average abundance of all the taxa in the same sample group. Taxa ranked below the top 10 are marked as other. Percentage of the most abundant taxon (blue) and of the taxa ranked below the 10 most abundant taxa (gray) are shown in the graph.

## Discussion

Fungal endophytes are recruited from the environment or transmitted endogenously through seeds. The environmentally-recruited taxa are assumed to be sporadic since they depend on their presence in the plant’s immediate surroundings, while taxa that persist within the plant are expected to be more stable, and therefore they might be regarded as core taxa (Christian et al. 2017). Importantly, many species can originate from both sources. The source of the taxa can greatly affect the composition, and therefore the functionality of FECs. In addition to the source of taxa, the composition of FECs is affected by environmental conditions and plant species, and changes during plant development and between plant organs, which complicates the discrimination between taxa from an external and endogenous origin.

To address the question of the transmission mode of fungal endophytes and its effect on FEC composition, we studied the correlations between FECs in the seeds and stems of three cereal species under natural conditions and in a controlled environment. We found that under natural conditions the seed population represented a subset of the stem population, suggesting that the seed population arises primarily from endogenous transmission. However, when we conducted similar experiments in a greenhouse, we found that the new seed FECs were as rich as the stems, and both the stems and new seeds contained numerous unique taxa that were not detected in the original seeds or in the soil. These results confirmed the notion that many taxa are vertically transmitted. However, they also revealed a significant influence of environmental conditions, not only on externally recruited taxa but also on the presence and levels of internally transmitted taxa.

Özkurt et al (2020) studied FECs in wheat and wild wheat and did not observe statistical differences in diversity between seed FECs of these plant species. However, the small sample size (58 individual seeds) and long duration of storage of wild wheat seeds might have contributed to the overall low diversity observed. Here, we collected samples from a total of eight sites with 40 samples for each plant species per location and analyzed a total of 420 seed samples. This robust analysis allowed us to capture a significant portion of the natural diversity. We showed that while the seed FECs in the three plant species contained similar taxa, the communities were nevertheless statistically different from one another. In particular, the two wild species were more diverse than the wheat FECs, as was previously found for stem FECs from *Ta* and *Td* (Sun et al. 2020b). The higher diversity of the wild plant FECs may reflect the more diverse environments in which they grow, as well as their higher genetic diversity compared with cultivated wheat (Sun et al. 2020b). *A. infectoria* dominated the seed FECs, raising the possibility that it represents an important core taxon in these plants and that it might be vertically transmitted. Fungi belonging to the genus *Alternaria*, including *A. infectoria*, were previously reported in wild and domesticated wheat (Ofek-Lalzar et al. 2016), as well as in seeds of other plant species, including rice (*Oryza sativa*) (Eyre et al. 2019), common oak (*Quercus robur* L.) (Abdelfattah et al. 2021), rapeseed (*Brassica napus*) (Morales Moreira et al. 2021) and radish (*Raphanus sativus*) (Rezki et al. 2018).

To further determine which taxa in the seeds are endogenously transmitted, we compared the seed and stem FECs. Our analyses showed a drastic decrease in the diversity and richness of seed FECs compared with the communities from stems; on average, the number of taxa in stems was about three times higher than in seeds and the stem FECs were twice as diverse. These results are in agreement with previous studies showing that seeds contain a lower number of fungal and bacterial species relative to other plant tissues, as well as to the environment (Newcombe et al. 2018; Hodgson et al. 2014; Chesneau et al. 2020; Ofek-Lalzar et al. 2016). Importantly, while the number of taxa in the seeds was greatly reduced compared with the stems, we found that close to 90% of the seed taxa (55 out of 62) were also found in the stems, suggesting that: (i) most of the seed taxa originated in the stem and are therefore endogenously transmitted, and (ii) most of the stem taxa do not reach the seeds. *Tremellomycetes* and *Saccharomycetes* yeast species were among the most highly abundant stem taxa that were not detected in seeds. Barret et al., 2015 found that during seed emergence, there is a reduction in bacterial and fungal diversity and attributed this reduction to an increase in fast growing taxa. Similarly, Hertz et al., 2016 showed that *Sporobolomyces roseus* was dominant in wheat during the first week of spike emergence and its levels decreased in later stages, concomitant with increased levels of *A. infectoria*. These studies together with our results support the possibility that the lower mobility of yeasts compared with filamentous species affects their ability to reach the seeds and establish themselves in the new community.

The high overlap of the seed taxa with stem taxa suggested that most of the seed-borne taxa are vertically transmitted (Fig 3A). However, the seeds and stems were collected in natural habitats, and therefore contribution from an external source such as pollen or direct seed infection cannot be excluded. For this reason, experiments aiming to isolate the infection source were carried out in controlled conditions. Such studies have usually focused on the connection between seed and roots or leaves, and therefore plants were sampled at the seedling stage (Abdelfattah et al. 2021; Özkurt et al. 2020; Barret et al. 2015; Hodgson et al. 2014). Here we tested the transmission of endophytes from seeds to stems to new seeds, and therefore grew plants to maturity during a period of five months. Despite autoclaving the soil twice, this measure could not completely prevent a buildup of soil population during the experiment, which probably contributed to the presence of some of the taxa found in the stems and new seeds population, especially at late sampling time points (Fig 5A, supplementary table S15). Nevertheless, aerial infection of seeds was minimized by covering the spikes before anthesis. Overall, despite a lack of complete sterility, the greenhouse experiments were performed in a relatively clean environment with tight control of the source of the endophytes, and prevention of aerial seed infection.

Much to our surprise, the stems, and new seeds from greenhouse-grown plants contained rich communities with a much higher number of taxa than either the soil or the original seeds, including numerous tissue-specific (either stem or new seed) taxa (Fig. 5A). Additionally, the FECs in the new seeds were as rich and diverse as the stem communities, unlike the highly reduced number of taxa observed in field-collected seeds. In a similar experiment, Latz et al., 2021 grew wheat plants both indoors and outdoors, studying progression of fungi from seeds to plant to seeds and how different potential sources of inoculum contribute to FECs. They showed that as much as 80% of the ASVs that were detected in plant tissues (seed stock, new seed, leaf, and root) could not be detected in the environment (air and soil) and concluded that airborne fungi are probably the primary inoculum source for fungal communities in aerial plant parts, whereas vertical transmission is likely to be insignificant.

Since the soil and original seeds were the only source for fungi in our experiment, our results argue for hidden taxa not revealed by the current sequencing procedure. It is possible that the high abundance of *A. infectoria* in the original seeds masked low abundant taxa. In stems and new seeds that did not contain a single highly dominant taxon, these rare taxa may have become visible because they could proliferate, as well as due to the greater sequencing depth. Indeed, 34% of the unique taxa that were discovered in the stems and seeds of greenhouse-grown plants were also found in field-collected plant samples and might have been present, yet undetected, in the original seeds. Regardless of the source of the new taxa, these results reveal substantial differences between the FECs in field and greenhouse-grown plants. Apart from the richer spectrum of taxa and their distribution, there is a drastic change in relationships between the FECs within different plant organs, suggesting changes in endogenous transmission. While we do not yet know what caused these changes, our findings emphasize the high sensitivity to external conditions of community assembly processes and transmission of endophytes within the plant.

Despite the differences between field-collected and greenhouse-grown samples, our results support endogenous transmission of a range of taxa. First, 16 taxa were found in all the tissues of greenhouse-grown plants (seeds, stems and new seeds), indicating seed-to-seed transmission. In addition, 51 taxa were found in the stems and new seeds of the greenhouse-grown plants as well as in seeds and stems collected from the field; it can be assumed that these taxa were present in the original seeds used in the greenhouse experiments, but below detection level, and were re-discovered when transmitted to the stems and new seeds. While taxa capable of endogenous transmission seem widespread, we observed significant differences in the relative importance of this transmission mode between fungal taxa. Most significantly, despite the high abundance of *A. infectoria*, our findings show that its main source in seeds is from aerial infection rather than endogenous transmission. This notion is supported by detection of airborne spores of *Alternaria* sp. including *A. infectoria* in the coastal plain of Israel (Waisel et al. 1997) and on pollen grains (Hodgson et al. 2014). Other taxa, such as *Cladosporium* sp. seem to depend mainly on endogenous transmission. The latter group probably represents the true core taxa and may therefore be of prime interest for future studies.

## Materials and methods

### Plant collections

Plant samples of bread wheat (*Triticum aestivum*; *Ta*), wild emmer wheat (*Triticum dicoccoides*; *Td*) and wild barley (*Hordeum spontaneum*; *Hs*) were collected from eight sites across Israel (Supplementary Table S1). For each plant species, we collected 40 spikes from each of four sites (160 spikes per plant species) in late spring and early summer of 2018. *Hs* stem samples were collected as described by (Sun et al., 2020a). *Ta* and *Td* stem samples were collected as described by (Sun et al., 2020b).

### Greenhouse experiments

For each of the three plant species we selected 15 seeds, each from a different plant. Seeds were surface sterilized and germinated as described by (Llorens et al., 2019). Germinated seeds were sown in 3-liter pots filled with double autoclaved peat soil and grown in the greenhouse with temperatures between 18°C and 24°C and a 14-hour light-10-hour dark regime. One tiller from each plant was collected after heading, which is the same developmental stage as the field-collected samples. To prevent aerial infection, spikes were covered with autoclaved paper bags at the beginning of anthesis. After harvest, the seeds were manually separated from the spikes and stored in paper bags. Five pots containing double autoclaved soil without plants were used to monitor the soil FECs at three time points: before sowing the seeds, at stem sampling, and dry seeds at the end of the experiment.

### Sample processing

#### Seed and stem samples

From each spike we sampled 2-3 seeds (100-150 mg). Seeds were surface sterilized by soaking in 3% commercial bleach for three minutes with agitation at 150 rpm, followed by three washes in sterilized double distilled water (DDW). Stems were sterilized as described in (Llorens et al. 2019) and sectioned into segments of 100-150mg. The sterilized seeds and stem samples were placed in 2 ml sterile tubes, flash frozen in liquid nitrogen and stored at -80°C.

#### Soil samples

Samples were taken from the middle of each pot, placed in a 50 ml tube, flash frozen in liquid nitrogen, and stored at -80°C until use.

### DNA extraction

#### Plant samples

DNA from seed and stems was extracted using a modified CTAB method according to (Sun et al. 2020a). In brief, samples were lyophilized overnight and ground to a powder using a Geno/Grinder 2000 (OPS Diagnostics, Bridgewater, NJ). 1 ml CTAB solution supplemented with 50 mg/ml proteinase-K and 1μl 2-mercaptoethanol were then added to each sample. The samples were incubated at 65°C for 45 minutes followed by 10 minutes at 95°C. After the samples cooled to room temperature, the DNA was separated with two rounds of chloroform-isoamylalcohol and ice-cold isopropanol precipitation overnight at -20°C. The DNA was washed twice with 70% ice-cold for 5 minutes at 4°C and 14,000 rpm. The DNA was resuspended with Tris-EDTA and 5% Chelex-100 chelating resin (Bio-Rad Laboratories, Hercules, CA).

#### Soil samples

DNA from soil samples was extracted using the DNeasy® PowerSoil® kit (QIAGEN, Hilden, Germany) according to (Berg-Lyons et al. 2018) with the following adjustments: the protocol was performed in individual tubes according to the manufacturer’s manual, C1 solution was supplemented with 10 mg/ml proteinase-K and pre-heated to 65°C before it was added to each sample, and all centrifuge stages were conducted at 14,000 x *g*.

### Amplicon sequencing

Amplicon library preparation and sequencing were performed by the Environmental Sample Preparation and Sequencing Facility (ESPSF) at the Argonne National Laboratory as described by (Sun et al. 2020a). ITS1f (5′-CTTGGTCATTTAGAGGAAGTAA-3′) and ITS2r (5′-GCTGCGTTCTTCATCGATGC-3′) primers were used to create amplicon libraries targeting the fungal ribosomal internal transcribed spacer 1 (ITS1) region (Walters et al. 2016). PCR reactions for plant samples were supplemented with a custom peptide nucleic acid (PNA) blocker (ITS1PNABlk: 50-O-E–E-GTCGTGTGGATTAAA-30; PNA BIO INC, Thousand Oaks, CA) to block host plant ITS sequences. In soil samples PCR PNA blocker was not used. Sequencing was done on the Illumina MiSeqsequencer (Illumina, San Diego, CA) on 251bp x 12bp x 251bp MiSeq run.

Field-collected stem sequences are available at NCBI Small Read Archive (SRA), *Ta* and *Td* BioProject ID: PRJNA509176, *Hs* BioProject ID: PRJNA592195. Field collected seed sequences are available at BioProject ID: PRJNA887576. All sequences generated at the greenhouse experiment are available at BioProject ID: PRJNA887702.

### Sequence processing

Field and greenhouse samples were processed and analyzed separately.

Reverse and forward reads were demultiplexed and a quality control test was performed in the QIIME2 platform (Bolyen et al. 2019). The DADA2 pipeline was used for sequence filtering, dereplication and taxonomical assignment (Callahan et al. 2016). Primers were trimmed using “Cutadapt”(Martin 2011). In the process, only forward reads were used due to the lower quality of the reverse reads. Taxonomic assignments of amplicon sequence variants (ASVs) were performed using UNITE fungal reference (Nilsson et al. 2019) and UNITE general FASTA release for eukaryotes to accurately filter non fungal ASVs. The ASVs were aligned against both databases using the Naïve Bayes approach with minimum 50 bootstrap. The final fungal set of ASVs was searched against the lab sequences database, which includes 700 bp amplicons of the ITS 4 region (Ofek-Lalzar et al. 2016) using BLAST, to increase the strength of the taxonomic assignment of the ASVs. Using the R package “phyloseq” (McMurdie and Holmes 2013), ASVs were agglomerated at the species level, and filtered for rare taxa. Singletons, doubletons, and samples with less than 1,000 sequences were removed from the dataset, then taxa with relative abundance <0.005 (0.5%) were also removed. Additionally, taxa with incidence <1% in field samples were removed.

### Data analysis

Data analysis was conducted in an R version 4.0.2 environment. The final set of reads was transformed using Hellinger transformation (Legendre and Gallagher 2001) for abundance analysis. Alpha diversity, Shannon Diversity Index and Observed taxa were calculated using “phyloseq”. Statistical analysis was conducted using “rstatix” in the R package (Alboukadel Kassambara 2021). Statistics for field samples were tested using Wilcoxon signed-rank test, and for greenhouse samples using Kruskal–Wallis one-way analysis of variance followed by pairwise Wilcoxon signed-rank test. Taxa differential abundance was tested by modifying the loop t-test described in (Wallen 2021) to a Wilcoxon test or Kruskal–Wallis test. *P* values generated in the analysis were adjusted using the Benjamini and Hochberg method (Benjamini and Hochberg 1995). Beta diversity was calculated with principal coordinates analysis (PCoA) using Bray–Curits in “phyloseq”. Permutational analysis of variance (PERMANOVA) was calculated using the *adonis* function in the “vegan” R package (Oksanen et al., 2020). To test statistical differences between multilevel pairwise comparison, the *pairwise*.*adonis* function was used (Martinez Arbizu 2020). Graphs were generated using “ggplot2” (Hadley Wickham 2016). Venn diagrams were generated using “ggvenn” package (Linlin Yan 2021). The heatmap was created using the “ampvis2” package (Andersen et al. 2018).

